# SARS-CoV-2 receptor and entry genes are expressed by sustentacular cells in the human olfactory neuroepithelium

**DOI:** 10.1101/2020.03.31.013268

**Authors:** Leon Fodoulian, Joel Tuberosa, Daniel Rossier, Madlaina Boillat, Chenda Kan, Véronique Pauli, Kristof Egervari, Johannes A. Lobrinus, Basile N. Landis, Alan Carleton, Ivan Rodriguez

## Abstract

Various reports indicate an association between COVID-19 and anosmia, suggesting an infection of the olfactory sensory epithelium, and thus a possible direct virus access to the brain. To test this hypothesis, we generated RNA-seq libraries from human olfactory neuroepithelia, in which we found substantial expression of the genes coding for the virus receptor angiotensin-converting enzyme-2 (ACE2), and for the virus internalization enhancer TMPRSS2. We analyzed a human olfactory single-cell RNA-seq dataset and determined that sustentacular cells, which maintain the integrity of olfactory sensory neurons, express *ACE2* and *TMPRSS2*. We then observed that the ACE2 protein was highly expressed in a subset of sustentacular cells in human and mouse olfactory tissues. Finally, we found *ACE2* transcripts in specific brain cell types, both in mice and humans. Sustentacular cells thus represent a potential entry door for SARS-CoV-2 in a neuronal sensory system that is in direct connection with the brain.

## Introduction

A novel virus from the Coronaviridae family, termed SARS-CoV-2, which emerged in December 2019 in East Asia, is currently expanding on the planet. Its exact history is unknown, but its genomic sequence suggests that it was transmitted from bats to humans via an intermediate animal host^1^. It is now transmitted from human to human^2^, and from human to cat^3^. Infection by SARS-CoV-2 is associated in our species with a severe respiratory syndrome called COVID-19, and is characterized by a substantial mortality rate^1,4,5^.

Entry of SARS-CoV-2 into target cells depends on the Spike protein (S), which is present on the virus capsid^6^. Viral attachment involves an interaction between S and the angiotensin-converting enzyme-2 (ACE2), located on the surface of the target cell. One reported mechanism facilitating virus entry consists, in association with ACE2, in the priming of S by the cellular serine protease TMPRSS2, also attached to the cellular membrane, which eventually leads to the fusion between the cellular and the viral membranes^1,7^. Expectedly, the main targets of SARS-CoV-2, respiratory cells that line the respiratory airways, coexpress *ACE2* and *TMPRSS2*^8^. Other proteins have been proposed to mediate SARS-CoV-2 internalization, in particular CD147^9^ as an alternative receptor, and cathepsin^10^ and furin-like proteases^6,11^ as internalization activators.

Starting with anecdotal reports, both from SARS-CoV-2-infected patients and medical staff which suggested an association between viral infection and alterations of olfactory perception^12-14^, the link between SARS-CoV-2 infection and olfactory dysfunction is today clearly established^15-20^. A smartphone app recording self-reported olfactory symptoms is even used to predict potential COVID-19^21^. The olfactory perturbations, which range from mild to more severe, are usually reversible, and affect up to 95% of COVID-19 patients depending on the report. Whether this olfactory perturbation results from a deterioration of the nasal sensor or of a more central perturbation is unclear, but the second hypothesis is not to be discarded since neurological manifestations of brain origin appear also to be associated with COVID-19 infection^22–26^.

The mammalian nasal cavity can be divided into two areas: the respiratory and the olfactory areas, that are anatomically, cellularly, and functionally different^27^ (Figure 1A). In humans, the respiratory part covers the major part of the nasal cavity. It includes the turbinates and is lined with a ciliated pseudostratified columnar epithelium. Its function is to humidify, cool or warm the incoming air, and to trap small particles before they get into the lungs. The second nasal area corresponds to the olfactory neuroepithelium. In our species, it is located in the very dorsal part of the cavity. There, it contacts volatile chemicals entering the nose, an interaction which represents the first step in the process that leads to the identification of a given smell. This epithelium is pseudostratified, and includes Bowman’s glands, olfactory sensory neurons, sustentacular cells, microvillar cells, globose and horizontal basal cells (cells that keep dividing during adult life and replenish the pool of sensory neurons)^28^. Each sensory neuron extends a single axon towards the olfactory bulb, that crosses the cribriform plate before reaching the olfactory bulb in the brain. On its apical side, the olfactory neuron extends a dendrite, which ends in multiple and long specialized cilia in contact with the outside world. These are covered with odorant receptors and bath in the mucus that lines the nasal cavity. Olfactory sensory neuron dendrites are enwrapped inside specialized cells, termed sustentacular cells^29^, whose nuclei and cell bodies line the external layer of the thick neuroepithelium (although they remain attached to the basal lamina). The role played by the latter in maintaining the integrity and function of the neuroepithelium is critical, in a very similar way that Sertoli cells support germ cell development and survival. Indeed, contact of the olfactory mucosa with various drugs (such as 3-methylindole^30^, the anti-thyroid drug methimazole^31^, or nickel sulfate^32^) to which sustentacular cells are very sensitive, leads to transient anosmia.

**Figure 1.**
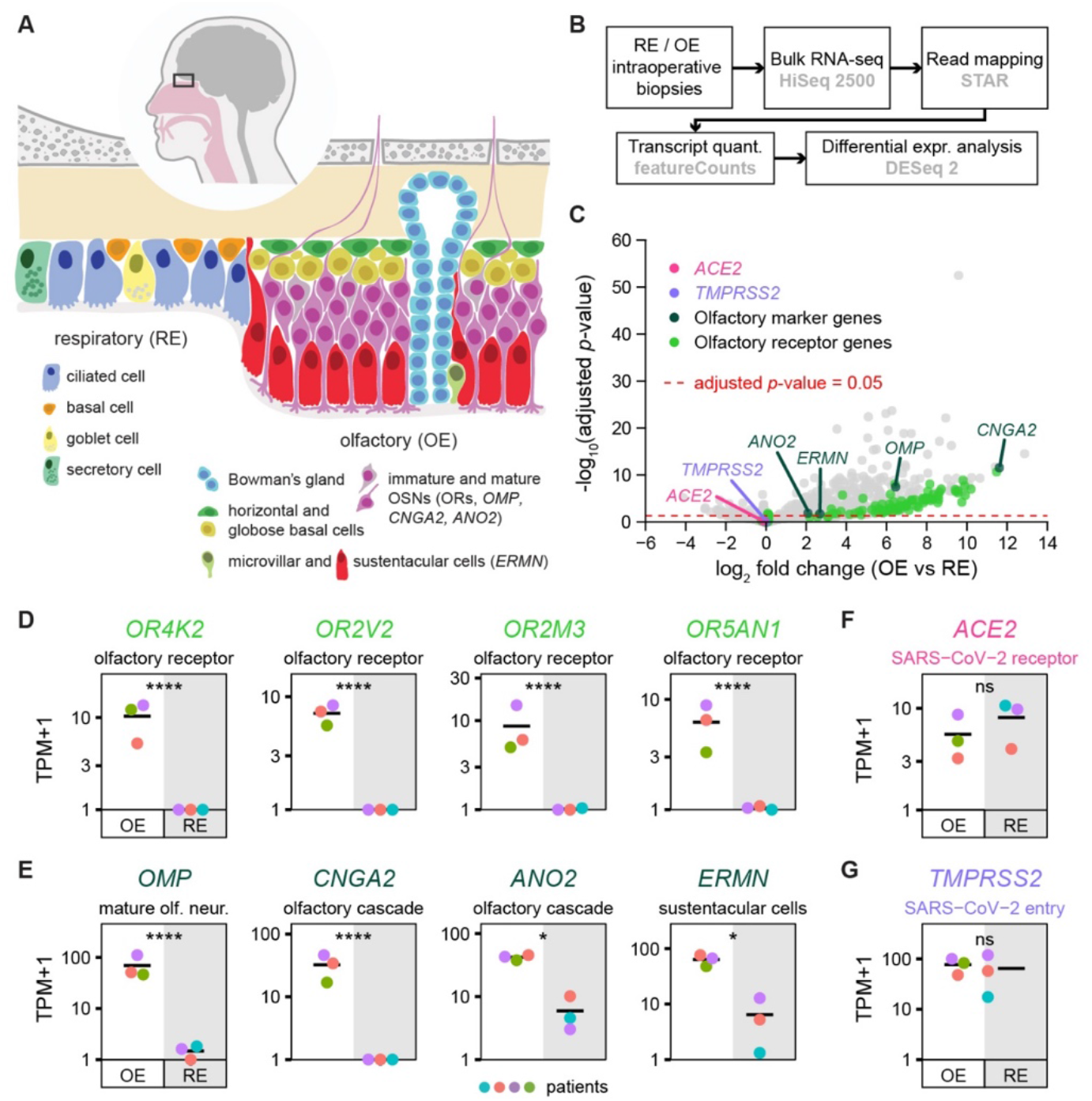
*ACE2* and *TMPRSS2* expression in olfactory and respiratory nasal epithelia. (**A**) Schematic of the human nasal respiratory and olfactory epithelia, with their respective cell types. (**B**) Design of the bulk tissue transcriptome analysis. OE biopsies: n=3, RE biopsies: n=3. (**C**) Volcano plot corresponding to the differential expression analysis, displaying the expression fold change and the associated adjusted *p*-value between olfactory and respiratory epithelia for each gene. OR genes, olfactory transduction cascade specific genes and SARS-CoV-2 entry genes *ACE2* and *TMPRSS2* are highlighted with a color code. (**D-G**) Transcript quantifications corresponding to (**D**) OR genes, (**E**) olfactory sensory epithelium markers and (**F** and **G**) SARS-CoV-2 entry proteins in olfactory and respiratory biopsies. A different color was attributed to the data of each patient (n=4). Olfactory and respiratory epithelium sample data points are shown on a white and grey background respectively. OE: olfactory epithelium, RE: respiratory epithelium.

Whether the apparent olfactory dysfunction associated with SARS-CoV-2 infection results from a general inflammation of the nasal cavity or from a more direct perturbation of the olfactory neuroepithelium is unknown. In any case, it is critical to determine whether this virus disposes of a niche to replicate just under the cribriform plate of the ethmoid bone, a structure with large holes through which olfactory neuron axonal projections directly contact the olfactory bulb, and offer a gateway to the brain.

We asked whether specific cells present in the human olfactory neuroepithelium as well as cells in the brain may represent targets to SARS-CoV-2, by looking at the molecular players involved in infection, both at the RNA and protein levels.

## Results

With the aim of exploring the potential expression of *ACE2* and *TMPRSS2* in olfactory sensory tissue, we collected biopsies via nasal endoscopic surgery from 4 adult patients. Samples of both nasal respiratory and olfactory sensory epithelia were harvested. Bulk tissue RNA was extracted, libraries generated, and sequenced (Figure 1B).

To assess the specificity of our dissection, we performed a differential expression analysis between the respiratory and olfactory epithelia datasets (Figure 1B). As expected, olfactory-specific genes, including olfactory receptor genes, *CNGA2* and *ANO2, OMP*, and *ERMN* (encoding critical elements of the olfactory transduction cascade, a specific marker of mature olfactory sensory neurons and a marker of sustentacular cells, respectively), were significantly enriched in olfactory samples (Figure 1C-E). The presence of *ACE2* and *TMPRSS2* transcripts was then evaluated. We observed a mean of 7.1 and 63.7 TPMs in the respiratory epithelium samples for *ACE2* and *TMPRSS2* respectively (Figure 1F,G), reflecting the presence of ciliated cells which represent targets of SARS-CoV-2. We found a mean of 4.6 and 76.1 TPMs in the sensory neuroepithelium samples for *ACE2* and *TMPRSS2* respectively (Figure 1F,G), indicating the presence in this tissue of cells that may express both genes, or of a mix of cells that express either *TMPRSS2* or *ACE2*.

To identify putative viral targets transcribing both *ACE2* and *TMPRSS2* and in the neuroepithelium, we took advantage of a very recently published dataset reported by Durante et al^33^. This dataset contains the transcriptome of 28’726 single cells, collected during nasal endoscopic surgery of 4 adult patients. Prior to any cell type analysis, we monitored the existence of cells that would transcribe both *ACE2* and *TMPRSS2* (Figure 2A). We then performed an aggregate analysis of the 28’726 single cells, and generated a UMAP dimensionality reduction plot, which allowed to display the clustering of the 26 different cell types reported in the original publication^33^, including olfactory, respiratory and immune cells (Figure 2B). We then identified all cells expressing *ACE2, TMPRSS2, ERMN* (an olfactory sustentacular cell marker) *and GSTA2* (a marker of respiratory ciliated and epithelial cells) (Figure 2C,E,H,I), and those coexpressing both *ACE2* and *TMPRSS2*, respectively (Figure 2D,H,I). Olfactory sensory neurons showed little or no expression of *TMPRSS2* and *ACE2*, respectively (Figure 2C). In contrast, different types of respiratory cells, but also olfactory sustentacular cells, expressed *ACE2*. Many of these cells also coexpressed *TMPRSS2*. To better evaluate the coexpression levels of *ACE2* and *TMPRSS2* in the different cellular populations, we plotted the mean normalized expression levels of *ACE2* and *TMPRSS2* corresponding to each of these subpopulations (Figure 2E). Sustentacular cells and ciliated respiratory cells exhibited the highest levels of expression of both genes. Populations containing olfactory horizontal cells, microvillar cells and Bowman’s glands also showed transcription of both genes, although at lower levels. To better evaluate the different expression levels inside the two cell types expressing the highest levels of both *ACE2* and *TMPRSS2*, we plotted the distribution of normalized transcript levels across all cells of both cell types (Figure 2F,G). Finally, the number (Figure 2H) and percentage (Figure 2I) of cells expressing *ACE2, TMPRSS* or both genes were evaluated.

**Figure 2.**
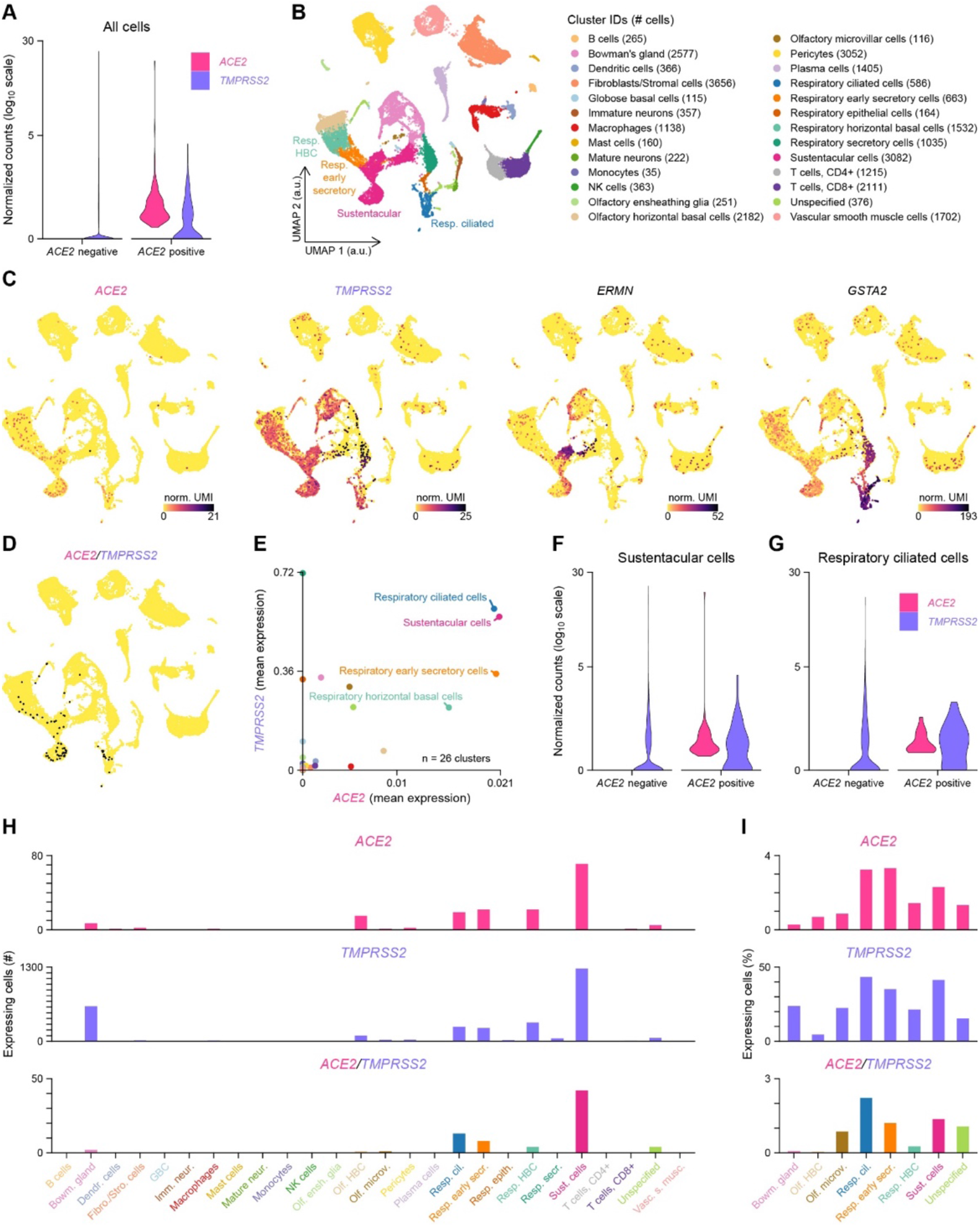
Coexpression of *ACE2* and *TMPRSS2* in sustentacular and respiratory ciliated cells. (**A**) Violin plots displaying *ACE2* and *TMPRSS2* normalized expression levels in all *ACE2* positive (n = 169) and *ACE2* negative (n = 28557) cells. (**B**) Visualization of the clustering results reported in Durante et al.^33^ on a uniform manifold approximation and projection (UMAP) plot. “Respiratory columnar cells” was renamed as “Respiratory early secretory cells” based on the expression of *SERPINB3*. a.u.: arbitrary units. (**C**) Normalized expression levels of *ACE2, TMPRSS2, ERMN* (a sustentacular cell gene marker) and *GSTA2* (a respiratory ciliated cell gene marker), shown on the UMAP plot. norm. UMI: normalized unique molecular identifier counts. (**D**) *ACE2* and *TMPRSS2* coexpressing cells, highlighted in black on the UMAP plot. (**E**) Coexpression of *ACE2* and *TMPRSS2* in the different cell clusters (n=26 clusters). Means of *ACE2* and *TMPRSS2* normalized expression levels are plotted. (**F,G**) Violin plots displaying *ACE2* and *TMPRSS2* normalized expression levels in (**F**) *ACE2* positive (n = 71) and *ACE2* negative (n = 3011) sustentacular cells, and in (**G**) *ACE2* positive (n = 19) and *ACE2* negative (n = 567) respiratory ciliated cells. (**H**) Number of cells per cluster expressing *ACE2* and/or *TMPRSS2*. (**I**) Percentage of cells per cluster expressing *ACE2* and/or *TMPRSS2* in all coexpressing clusters (i.e. clusters with double-positive cells). Color gradient scales in panel **C** and y-axis scales in panels **A**, **F** and **G** are in log_10_.

Respiratory cells are thus not the only cells in contact with the outside world that exhibit the molecular keys involved in SARS-CoV-2 entry in the nose. Sustentacular cells, which are localized at a porous boundary between the central nervous system and the olfactory cavity, share the same characteristics.

Quantification of gene transcripts at the single cell level, a commonly used proxy of protein abundance, often leads to very inaccurate predictions, up to a complete discordance between real transcript and protein levels^34,35^. Since it is the protein that matters here, and at the individual cell level, we investigated the expression of ACE2 and TMPRSS2 in tissues by immunohistochemistry. Given that ortholog tissuespecificity is highly conserved between tetrapod species^36^, and in particular between mouse and humans^37^, we analyzed mouse tissues in parallel to human ones.

We first evaluated the potential ACE2 immunoreactivity of the mouse olfactory epithelium (Figure 3A,B). To assess antibody specificity, we used independently three different anti-ACE2 antibodies, which were raised against the extracellular part (Ab1 and Ab3 developed in goat and rabbit respectively) or the intracellular part of ACE2 (Ab2, developed in rabbit). These antibodies were first evaluated to label colon and kidney tissues, which contain previously described ACE2-expressing cell types (Supplementary Figure 1A-J). In the olfactory cavity, a strong and defined labeling was observed in the respiratory epithelium in the nasal cavity and nasopharyngeal duct (Supplementary Figure 2A-H), and in a subset of sustentacular cells lining the nasal cavity (Figure 3C-G and Supplementary Figure 2B). These were almost exclusively located in the dorsal part of the epithelium, with a relatively abrupt transition area (Figure 3H,I). All antibodies showed a labeling of the apical portion of sustentacular cells, except for a slight labeling of their somata with Ab2 (Figure 3D). To clearly differentiate olfactory sensory neurons from sustentacular cells, we colabeled the sections with TUJ1, a marker of neurons, OMP, a marker of mature sensory neurons, and ERMN, a marker of sustentacular cells (Figure 3J-P). The localization of ACE2 was at the very luminal border of the sustentacular cells, sandwiched between the sensory neurons TUJ1-expressing cilia, and ERMN, which labels the apical part of sustentacular cells (Figure 3J-S). The whole epithelium, as expected based on the single-cell RNA-seq data, was positive for TMPRSS2 (Figure 3N).

**Figure 3.**
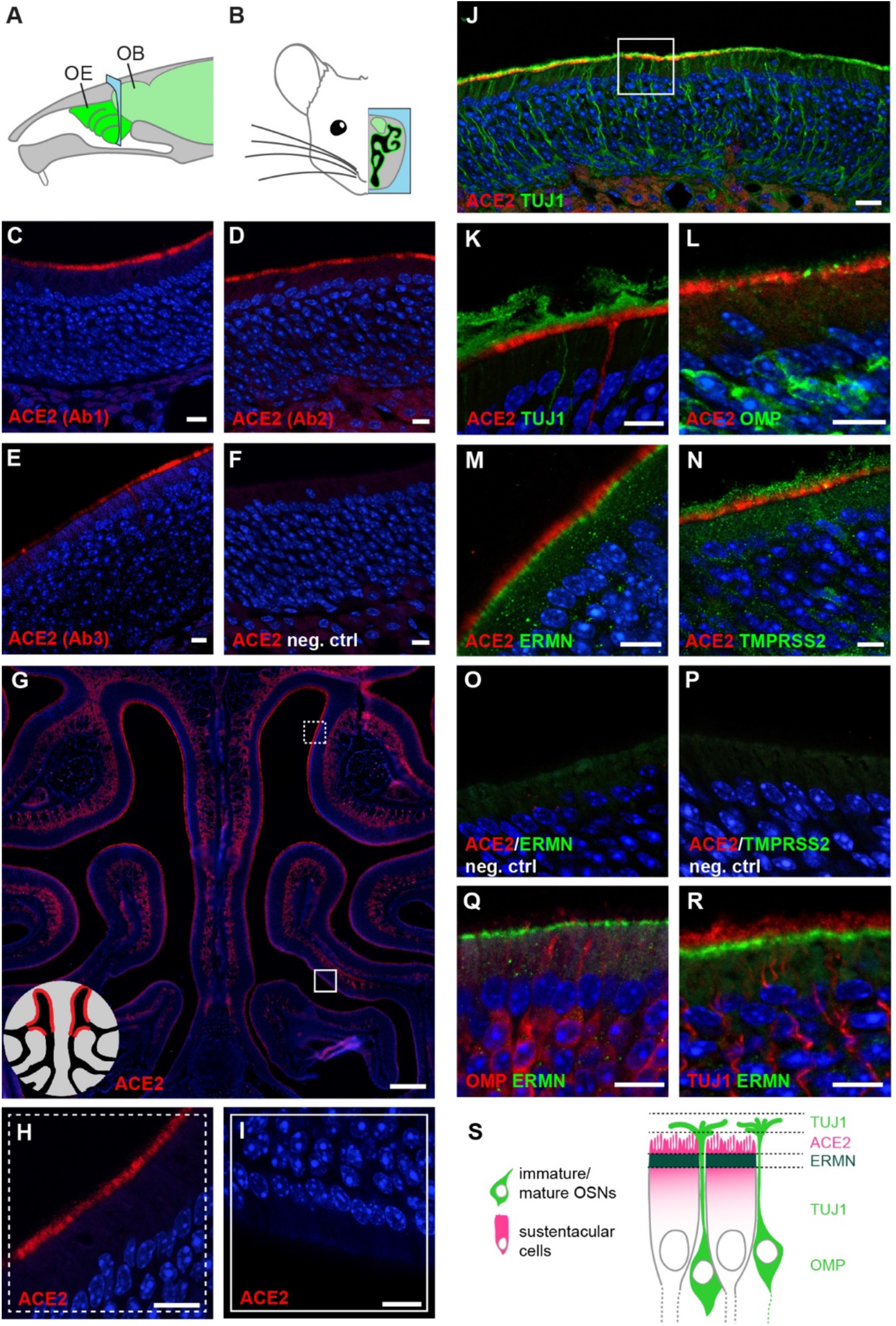
*Ace2* expression in the mouse olfactory neuroepithelium. (**A**) Schematic representation of the mouse olfactory system. The sensory tissue and the brain are highlighted in green. The blue frame indicates the coronal section level corresponding to the next panels. OB: olfactory bulb, OE: olfactory epithelium. (**B**) Schematic representation of a coronal section of the mouse olfactory system. The OE is highlighted in green, and the brain, including the olfactory bulb, in light green. (**C-F**) Immunostainings on coronal sections of mouse olfactory neuroepithelium with three different anti-ACE2 antibodies (red) and a negative control. (**C**) ACE2 Ab1, polyclonal antibody against the extracellular domain of mouse ACE2. (**D**) ACE2 Ab2, polyclonal antibody against the intracellular domain of human ACE2. (**E**) ACE2 Ab3, monoclonal antibody against the extracellular domain of human ACE2. (**F**) Negative control immunostaining without primary antibody against ACE2. (**G**) Immunostaining for ACE2 (red) on a coronal section of the mouse olfactory neuroepithelium. Scale bar: 0.2 mm. Squares indicate zones magnified in (**H**) and in (**I**). The schematic on the lower left indicates that ACE2 expression is observed in the dorsal (**H**), but not in the ventral part (**I**) of the olfactory epithelium. (**J**) Double immunostaining for ACE2 (red) and the neuronal marker TUJ1 (green) in the ACE2-positive part of the mouse olfactory neuroepithelium. The square indicates the portion of the MOE that is magnified in panels (**K-R**). Scale bar: 20 μm. (**K-N**) Double immunostaining for ACE2 (red) together with (**K**) the neuronal marker TUJ1 (green), (**L**) the marker of mature olfactory sensory neurons OMP (green), (**M**) the marker for sustentacular cells ERMN (green) and (**N**) TMPRSS2 (green). (**O-P**) Negative control immunostaining without primary antibody against (**O**) ERMN and (**P**) TMPRSS2. (**Q,R**) Double immunostaining for the sustentacular cell marker ERMN (red) together with (**Q**) OMP and (**R**) TUJ1. (**S**) Schematic of the external part of the olfactory epithelium, highlighting the relative positions of ACE2, TUJ1, ERMN and OMP according to the immunostainings shown in panels (**K**-**R**). TUJ1 is present in the dendrites, axons and somata of olfactory sensory neurons, OMP more concentrated in the cell body of olfactory sensory neurons, and ACE2 and ERMN in the apical portion of sustentacular cells. All sections with immunostaining were counterstained with DAPI (blue). When not indicated, scale bars are 10 μm.

We then evaluated the expression of ACE2 in the human nasal cavity by immunohistochemistry. Similarly to what is observed in the mouse, the human neuroepithelium can be discriminated from the respiratory epithelium by the presence of TUJ1-positive olfactory sensory neurons (Figure 4A), ERMN-positive sustentacular cells along the apical border of the tissue (Figure 4A, Supplementary Figure 3C-E), as well as the absence of alcian blue-labeled goblet cells (Figure 4B). In accordance with the single-cell data, a strong labeling of ACE2 was observed in some sustentacular cells of the sensory epithelium (Figure 4C-G). However, the very apical localization of ACE2 observed in the mouse neuroepithelium was not recapitulated in humans, but ACE2 was rather enriched in the apical half of the cells. A similar somatic staining was also observed, with the same antibody, in proximal tubule cells of the mouse kidney (Supplementary Figure 1D). Congruently with its pervasiveness in the single-cell transcriptomes, TMPRSS2 labeling was observed in many constituents of the nasal epithelium (Figure 4H).

**Figure 4.**
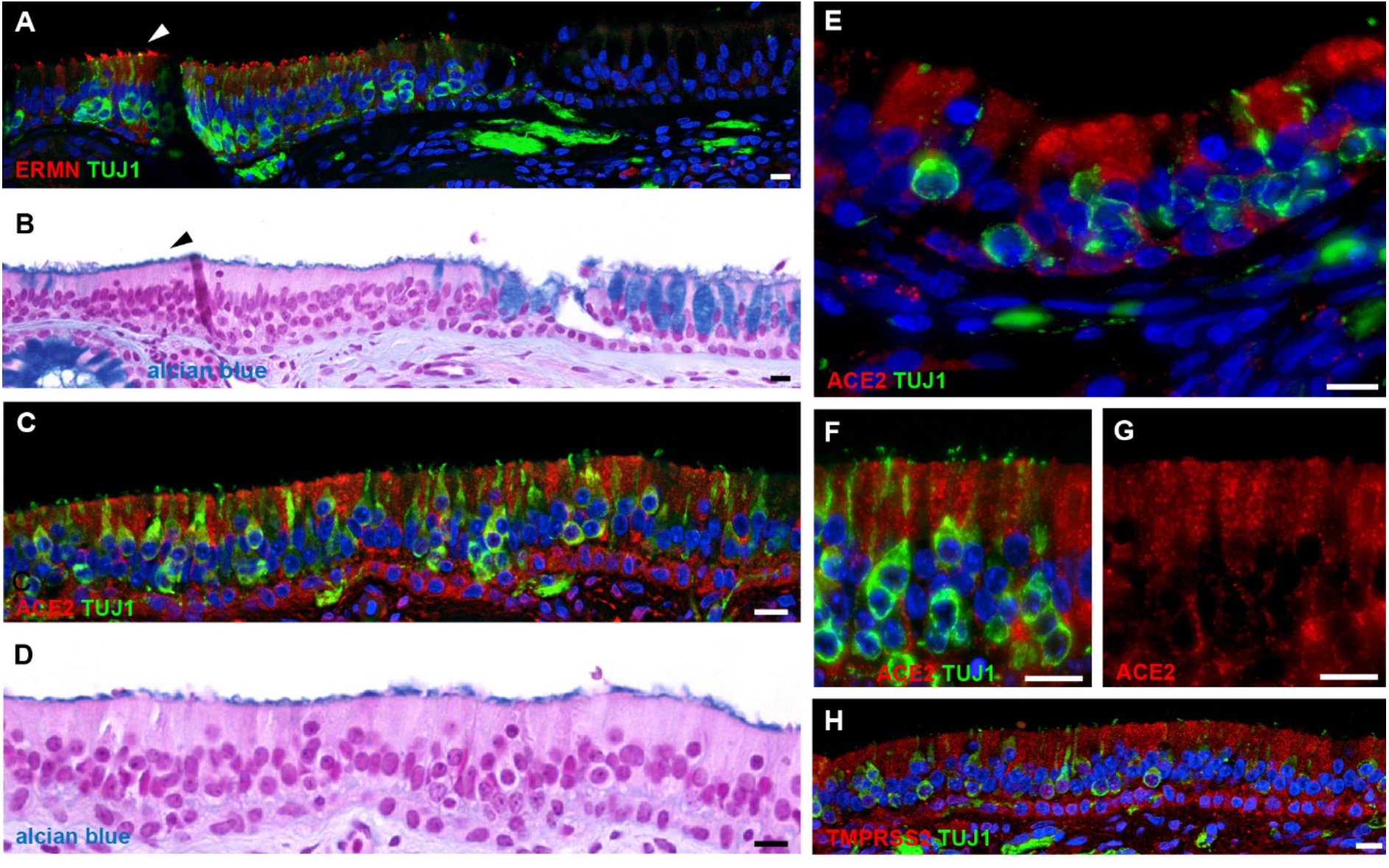
*ACE2* expression in the human olfactory neuroepithelium. (**A,B**) Section of human respiratory epithelium with (**A**) immunostaining for ERMN (red) and the neuronal marker TUJ1 (green), and (**B**), the same section stained with alcian blue and counterstained with nuclear fast red, highlighting a transition between sensory (left) and respiratory (right, note the goblet cells) epithelium. ERMN is expressed in the apical part of sustentacular cells within the olfactory neuroepithelium (arrowheads). (**C,E**) A portion of human olfactory neuroepithelium immunostained for ACE2 (red) and the neuronal marker TUJ1 (green), and (**D**) with alcian blue (note the absence of goblet cells). (**F,G**) Higher magnification of the external border of the human olfactory neuroepithelium. The same region is shown with (**F**) and without (**G**) TUJ1 staining. Nuclei were counterstained with DAPI (blue) or FastRed (red). Scale bars: 10 μm.

SARS-CoV (i.e. the virus responsible for the previous SARS epidemic in 2002), which genome is closely related to the one of SARS-CoV-2 and whose effects on human tissues appear similar to those observed with SARS-CoV-2, has been observed in human brains^38-40^. Could SARS-CoV-2 replicate in the olfactory epithelium and could the infection spread to the brain? To explore the potential receptivity to SARS-CoV-2 infection of the various cell types in the brain, we evaluated, again, the potential coexpression of *ACE2* and *TMPRSS2* in neuronal and non-neuronal cell types in the central nervous system of both mice and humans.

For the mouse, we took advantage of two available single-cell RNA-seq datasets^41,42^, consisting in two broad collections of mouse brain cell types. In the first collection, published by Zeisel et al^41^, that include non-neuronal cells, we found a very limited number of cell types that express *Ace2* (Figure 5A). These cells are related to pericytes and coexpress the mural cell marker *Rgs5*. They express *Ace2* transcripts, as well as *Rgs5*, a pericyte marker (Figure 5B). *Tmprss2*, whose transcripts were barely detected, was expressed by even less cell types (Figure 5A), with Purkinje cells displaying the highest expression level. These observations were confirmed by the second mouse dataset^42^, in which mural cells were the main cell type that expressed *Ace2* (Figure 5C). None of these cell populations coexpressed *Tmprss2*. Although showing substantial expression of ACE2 in the mouse brain, our data did not replicate the widespread expression of ACE2 in the central nervous system previously reported by others, which found ACE2 in the motor cortex, raphe, and in nuclei involved in the regulation of cardiovascular function^43^. To precisely evaluate the expression of ACE2 in the mouse brain, and in particular at the interface with the olfactory sensory neuroepithelium, we performed an immunostaining of mouse olfactory bulb sections with the three different anti-ACE2 antibodies. A strong labeling of capillary associated cells was observed through the different layers of the olfactory bulb (Figures 5D-H), in agreement with the single-cell RNA-seq observations.

**Figure 5.**
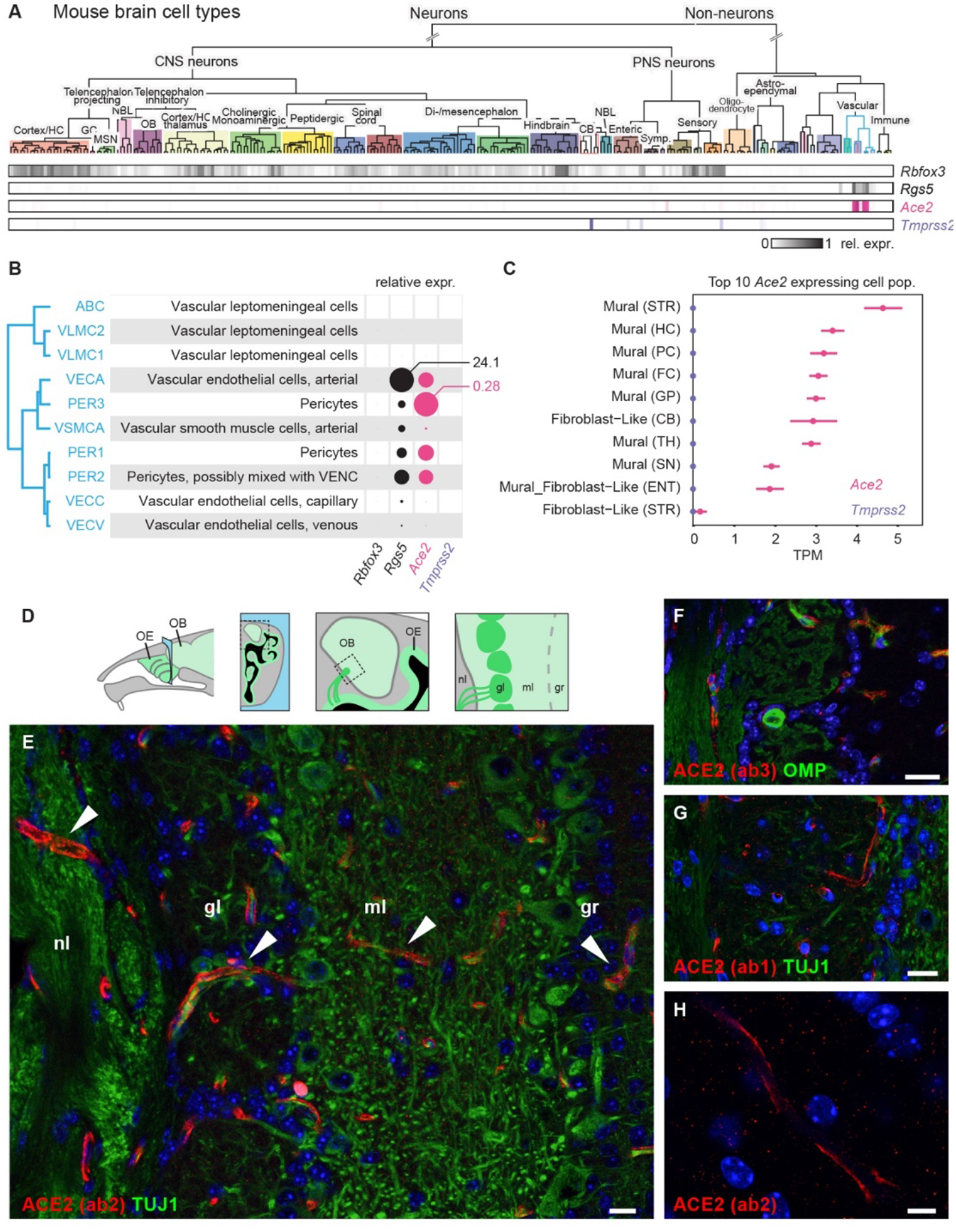
*Ace2* and *Tmprss2* expression in the mouse brain. (**A**) Mouse brain cell type classification from Zeisel et al.^41^ were visualized via http://mousebrain.org/. Rows with shaded boxes below the tree indicate the gene mean expression levels for each cell type, relative to its maximum. *Rbfox3*, encoding the neuronal marker NEUN, and *Rgs5*, a marker of pericytes, are shown in comparison to *Ace2* and *Tmprss2*. (**B**) Details of the gene expression levels in pericytes (blue edges clade of the classification dendrogram in (**A**)). Discs represent mean relative expression levels. Maximum values of mean normalized expression levels for *Rgs5* and *Ace2* are connected to the corresponding circles. (**C**) *Ace2* and *Tmprss2* expression levels in the top 10 *Ace2*-expressing cell populations from the http://dropviz.org/ dataset^42^. Abbreviations in parentheses denote the origin brain structure. STR: striatum, HC: hippocampus, PC: posterior cortex, FC: frontal cortex, GP: globus pallidus, CB: cerebellum, TH: thalamus, SN: substantia nigra, ENT: entopeduncular. (**D**) Similar to Figure 3, subsequent panels show coronal sections (blue area) crossing the nasal cavity and the olfactory bulb. As represented in this schematic, olfactory sensory neurons from the olfactory epithelium send axonal projections to the olfactory bulb through the cribriform plate of the ethmoid bone, along the nerve layer. OB: olfactory bulb, OE: olfactory epithelium, nl: nerve layer, gl: glomerular layer, ml: mitral cell layer, gr: granule cell layer. (**E-G**) Immunostaining for ACE2 (red) and for the neuronal marker TUJ1 (green) on sections of the mouse olfactory bulb. White arroheads indicate the presence of ACE2 in the pericytes of capillaries. Scale bars: 20 μm. (H) Magnification of capillary ACE2 immunostaining (red). Scale bar: 5 μm. All sections were counterstained with DAPI (blue).

Finally, we explored two single-nucleus RNA-seq datasets to evaluate *ACE2* expression in the human brain. In the first dataset, published Lake et al.^44^, we found neuronal and glial cell types, in particular Purkinje neurons and cerebellar astrocytes, that expressed *ACE2* or *TMPRSS2* (Supplementary Figure 4A-E). In the second dataset, publicly available in the Allen Brain Map cell types database, we found expression of *ACE2* (although at relatively limited levels) in various non-neuronal and neuronal types, in particular astrocytes, oligodendrocytes, and neurons from the cortical layer 5 (Figure 6A-G), a portion of which coexpressed TMPRSS2 (Figure 6C-E). ACE2 and TMPRSS2 are therefore expressed in a number of cell types in the human brain.

**Figure 6.**
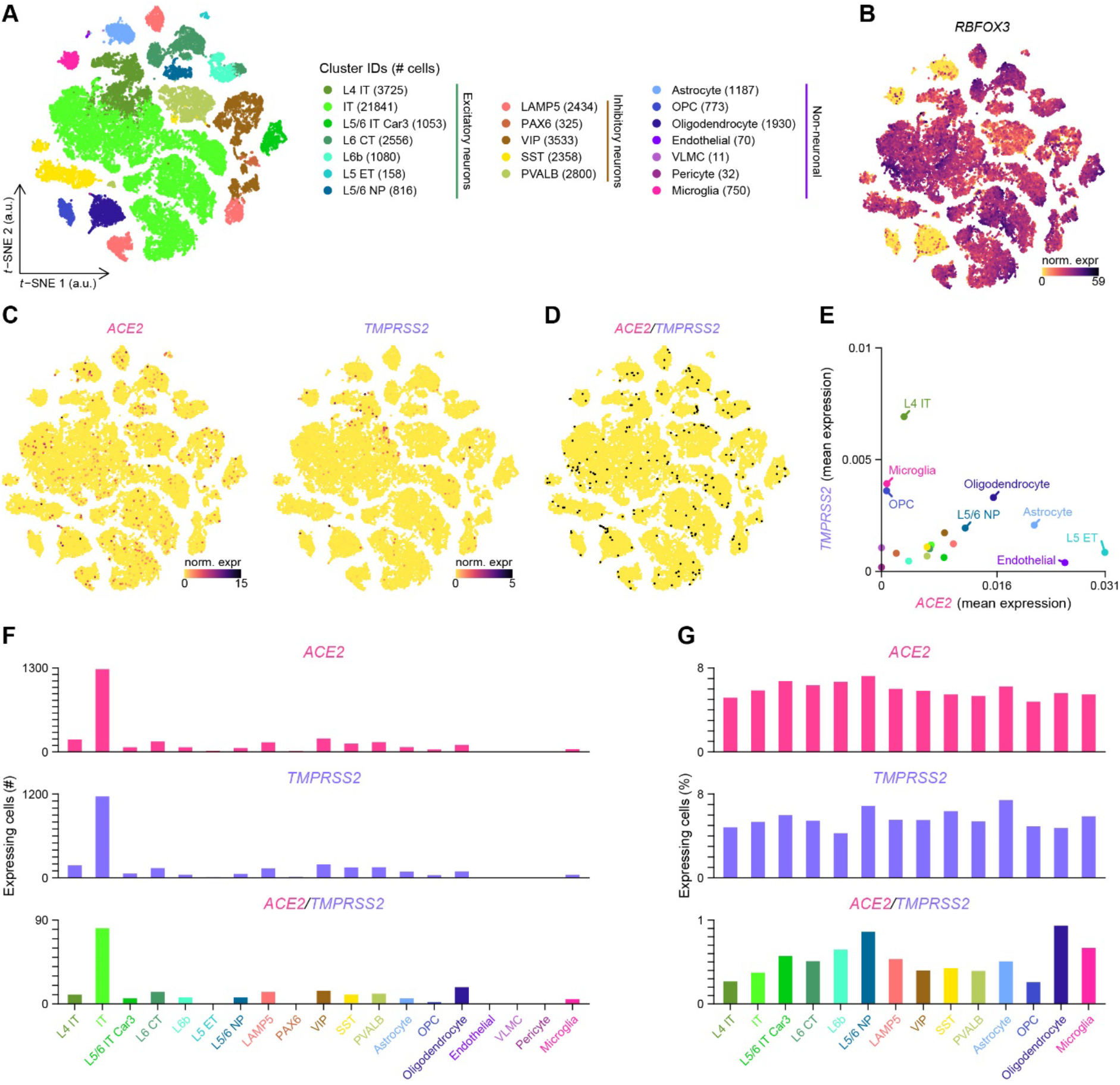
Coexpression of *ACE2* and *TMPRSS2* in the human brain. (**A**) Visualization of the human brain cell-type subclasses on a *t*-distributed stochastic neighbor embedding (*t*-SNE) plot. a.u.: arbitrary units. (**B-C**) Normalized expression levels of (**B**) *RBFOX3* (which encodes for the neuronal marker NEUN), (**C**) *ACE2* and *TMPRSS2*, shown on the *t*-SNE plot. norm.expr: normalized expression levels. (**D**) *ACE2* and *TMPRSS2* coexpressing cells, highlighted in black on the *t*-SNE plot. norm. expr:. (**E**) Coexpression of *ACE2* and *TMPRSS2* in the different subclasses (n=19 subclasses). Means of *ACE2* and *TMPRSS2* normalized expression levels are plotted. (**F**) Number of cells per subclass expressing *ACE2* and/or *TMPRSS2*. (**G**) Percentage of cells per subclass expressing *ACE2* and/or *TMPRSS2* in all coexpressing subclasses (i.e. subclasses with double-positive cells). Color gradient scales in panels **B** and **C** are in log_10_. L: cortical layer, IT: intratelencephalic, CT: corticothalamic, ET: extratelencephalic, NP: near-projecting, LAMP5: lysosomal-associated membrane protein family member 5, PAX6: paired box 6, VIP: vasoactive intestinal polypeptide, SST: somatostatin, PVALB: parvalbumin, OPC: oligodendrocyte precursor cell, VLMC: vascular leptomeningeal cell.

## Discussion

The massive and recent interest linked to COVID-19 has led to a number of reports, many of which are still at a preprint repository stage. A major question, addressed by some of these reports is the viral receptor distribution in the human airways^45-47^.

Taking a multidisciplinary approach, based both on our own and publicly available RNA-seq datasets, and on immunohistochemical stainings of mouse and human tissues, we demonstrate here that a subset of olfactory sustentacular cells in the olfactory neuroepithelium, but not olfactory sensory neurons, expresses *ACE2*, the main player in the binding and entry of the SARS-CoV-2 into human cells. These cells were found to coexpress *TMPRSS2*, a serine protease known to facilitate viral entry. We also found ACE2 expression in specific cell types of the brain, including neuronal and non-neuronal cell types.

We first determined, using transcriptomic analyses of whole tissue and single cells from human olfactory epithelia, that a subset of sustentacular cells expresses ACE2. We recently reported this finding as a preprint on BioRxiv, which is in agreement with another preprint on the same server^45^. We then confirmed this finding in mouse and human tissues by immunohistochemistry. In the mouse, in which the olfactory mucosa is particularly well organized both in terms of pseudostratified layers and in terms of its very strict separation from the respiratory epithelium, we observed a clear expression of ACE2 in the apical border of sustentacular cells (in agreement with a very recent report citing our BioRxiv article^46^). This distribution was however non-homogenous, since ACE2 was present in the most dorsally-located sustentacular cells, but was entirely absent from more ventrally-located ones. Such dichotomy between the dorsal and the ventral neuroepithelium is reminiscent, in the mouse, of the dorsally-located olfactory neurons, which are known to share molecular markers that are dissimilar from those present in more ventral neurons^48^. This known contrast is not limited to the neuronal identities, since the function and response to stress of these dorsally-located neurons also appears to be different^49,50^. We now add a nonneuronal cell type to this dorsoventral neural dichotomy. In the human olfactory neuroepithelium, a similar labeling of ACE2 was observed in sustentacular cells, although the staining was mainly present in the somata. Again, like in the mouse, some of these cells were positive and others were negative for ACE2. These human ACE2-expressing sustentacular cells were found isolated or in small clusters, intermingled with sustentacular cells negative for ACE2, or even sometimes intermingled with respiratory epithelium, an expected observation given the relatively poor organization of the human olfactory neuroepithelium, in particular in aged individuals.

How likely is it that the coexpression of *ACE2* in olfactory sustentacular cells is involved in the SARS-CoV-2-induced anosmia? And why are horizontal basal cells that also express *ACE2* not prime candidates? First, given the slow rate of neuronal renewal in the main olfactory epithelium (a few months) and the apparent rapid development of SARS-CoV-2-triggered anosmia, a perturbation of the stem cell pool constituted by the horizontal basal cells appears unlikely to represent an immediate disturbance to the system. Second, the amounts of *ACE2* transcripts as well as the intensity of the immunohistological stainings we observed in sustentacular cells are in the same range as the ones observed in respiratory ciliated cells, suggesting at least that sustentacular cells play in the same league in terms of SARS-CoV-2 receptors.

Third, sustentacular cells are in direct contact with the olfactory cavity, and thus interact with everything that enters the nose. Fourth, sustentacular cells play a critical role in the maintenance of the olfactory neuroepithelium integrity: their loss or suffering would cause disaggregation or malfunctioning of the neuroepithelium, respectively, leading to anosmia. Finally, given that the role played by ACE2 in sustentacular cells is unknown, the hijack of this receptor by SARS-COV-2 may by itself perturb the functioning of the olfactory sensor. Taken together, and despite the fact that one cannot exclude inflammation and the infection of other non-neuronal cell types in the olfactory neuroepithelium as an origin of the SARS-CoV-2-induced anosmia, the link between the viral molecular entry tools expressed by olfactory sustentacular cells and the SARS-CoV-2-induced chemosensory alteration appears quite credible.

The idea that viruses may affect directly or indirectly the integrity and function of the sensory part of the olfactory system is not new. Some viruses indeed disturb the neuroepithelium in different ways, and often alter specific cell types, including neurons. For example and among many others, olfactory sensory neurons and sustentacular cells were shown to be major portals of host entry of the Murid Herpesvirus-4 (MuHV-4)^51^. Again, the direct contact of this latter cell type with the respiratory tract, as well as its phagocytic activity, may explain its particular sensitivity to viral infection^52^. The cell type targeted by viruses matters, in particular to fight infection using drugs. For instance, in the case of a MuHV-4 challenge, olfactory sensory neurons are not responsive to interferon treatment (IFNγ), while sustentacular cells are^53^.

SARS-CoV-2-infection has not only been linked to the loss of a single chemosensory ability. Indeed, in addition to anosmia, ageusia has been reported^12^. Whether these reports truly reflect taste anomalies, or rather olfactory perturbations that may drastically affect the flavor of food, is unclear. However, this potential double effect on two chemosensory systems that share nothing at the periphery may suggest a more central alteration, involving for example a direct infection of the brain by SARS-CoV-2. But this would require potential targets. We identified various cell types in the brain that express ACE2, although at relatively low levels in humans. In the mouse, we detected ACE2 expression in brain vascular endothelial cells and pericytes, but not in neurons. In humans, we found *ACE2* transcripts in non-neuronal cell types such as astrocytes and microglia, but also in neurons, in particular Purkinje and cortical layer 5 neurons. Whether these different cell types reflect differences in tissue specificity between the two mammalian species is unclear, since RNA-seq discordances may results from differences in the sequencing depth, sequencing protocols (singlenucleus versus single-cell RNA-seq), dissimilar mRNA half-lives, or variable cell type selection strategies between the different studies. But whatever the reason for the discordances, these data point to potential targets for SARS-CoV-2 infection in brain cells.

Similar to the infection of the olfactory neuroepithelium by SARS-CoV-2, the transport of viral particles from the olfactory mucosa to the brain would not be a first. In 1935 already, the olfactory route was hypothesized (because of its direct connection with the brain), to represent a portal for entry of viruses into the central nervous system^54^. We know today that the olfactory nerve route is indeed used by various viruses to reach the brain, including HSV1, poliovirus, MHV (a coronavirus), paramyxoviruses, Hendra virus, Nipah virus, influenza virus, adenoviruses, bunyaviruses, VSV, and many others^55^. Furthermore, this direct olfactory path is so efficient that it is taken advantage of to facilitate access of drugs into the brain.

SARS-CoV particles have in fact been found in the human brain^38-40^. Adding to this idea, brain lesions were observed in a transgenic mouse model expressing the human *ACE2* in the nose and infected intranasally with SARS-CoV, brain lesions were observed^56^. Moreover, a retrospective case study on 214 COVID-19 patients reported neurological manifestations possibly correlated with the severity of the disease^57^ (with the confounding factor that old people are more likely to develop severe disease). As worrying are suggestions that a subset of patients infected with the virus but lacking respiratory symptoms may exhibit neurologic symptoms^22^, and that they may be associated with encelphalitis^23,25^, Miller Fisher syndrome and polyneuritis cranialis^26^. Given, as we showed here, the presence in the human brain of various neuronal and non-neuronal cell populations expressing ACE2, this certainly appears to be a line of investigation worth pursuing.

## Materials and Methods

### Human biopsies and bulk RNA sequencing

After authorization from the University of Geneva ethics commission, human biopsies were collected at the Geneva University Hospital. For whole tissue RNA-seq, samples of 4 individuals (3 males and 1 females) were resected during nasal cavity surgeries under general anesthesia. Biopsies consisted in small pieces of tissue of approximately 3 mm in diameter, from the respiratory or the sensory epithelia. Biopsies were snap frozen in liquid nitrogen immediately upon collection. For RNA extraction, tissues were placed in RLT Buffer (Qiagen) with beta-mercaptoethanol and lysed with stainless steel balls (5mm diameter, Schieritz and Huenstein AG) using a homogenizer (Big Prep, MB Biomedicals). Total RNA was isolated using RNeasy Mini Kit (Qiagen) following the manufacturer protocol. The quality and quantity of total RNA were evaluated with a Bioanalyzer (Agilent). Stranded cDNA libraries with suitable adapters for multiplexing were generated with Truseq RNA and DNA sample preparation kits (Illumina) following ribodepletion of the total RNA (200ng of total RNA per sample). Samples were multiplexed for sequencing in a HiSeq^®^2500 Sequencing system, generating 100bp single-end reads.

### Bulk RNA sequencing data analysis

RNA-seq reads were mapped onto the GRCh38 human genome assembly with STAR^58^ version 2.7.0a using the Ensembl v99 gene annotation file (GTF). Mutlimapped reads were filtered out with the option --outFilterMultimapNmax set to 1. Reversely stranded gene expression quantification was carried using featureCounts^59^ version 1.6.3 and a modified version of the aforementioned GTF file. The records corresponding to *AC097625.2*, a newly annotated long non-coding RNA that spans and shares most exons of *ACE2*, were removed from the GTF file as they hampered *ACE2* quantification (due to ambiguous read counting). This gene had a total of 3 detected counts in two samples and its removal from the GTF file led on average to a 9-fold increase in *ACE2* counts. Note that this annotation was absent from and did not affect the single-cell RNA-sequencing datasets as it first appeared in the GENCODE release 30 and the Ensembl release 96 of the GRCh38 GTF files. Differential gene expression analysis between main olfactory epithelial (MOE) biopsies and respiratory epithelial (RE) biopsies was performed using the DESeq2^60^ Bioconductor package version 1.22.2 in R version 3.5.0. After fitting a negative binomial generalized linear model (GLM), the Wald test (two-tailed) was used to test for significance of gene expression differences as a function of biopsies at a log_2_ fold change threshold of 0. To control the false discovery rate, the Wald test p-values were adjusted for multiple comparisons using the Benjamini–Hochberg procedure^61^. Patient identifications were not added to the model to control for potential differences in gene expression between individuals (i.e. batch effects) due to the underpowered experimental design highlighted by the 3 samples per condition. The Approximate Posterior Estimation for GLM coefficients (an adaptive Bayesian shrinkage estimator) implemented in the apeglm^62^ Bioconductor package version 1.4.2 was used to shrink log_2_ fold changes. During shrinkage, the maximum likelihood estimates (i.e. log_2_ fold changes) and their associated standard errors (both in log scale) were used to adapt the scale of the prior. The results of this analysis are represented as a volcano plot in Figure 1C. To display the expression levels of selected genes in individual samples (Figure 1D-G), TPM values were calculated for each gene within each sample to normalize for sequencing depth differences. Olfactory cascade genes (i.e. *OMP, CNGA2* and *ANO2*) and the top 4 most differentially expressed OR genes (with the lowest adjusted *p*-values) were selected as markers of the olfactory neurosensory epithelium. ERMN was selected as a marker of sustentacular cells^29,63^. All datasets generated during and/or analyzed during the current study are available on request.

### Single-cell RNA sequencing data analysis of human olfactory epithelial cells

#### Data download

The processed 10X Genomics output files of all four patients of the single-cell RNA-seq dataset reported in Durante et al.^33^ were downloaded from the NCBI GEO database with the accession number GSE139522. This dataset consists in human olfactory and respiratory epithelial cells collected from four patients aged between 41 and 52 years. The data was analyzed in R version 3.5.0 using the Seurat^64-66^ R package version 3.1.4 following the indications described in Durante et al.^33^ and using custom scripts.

#### Cell and gene filtering

The 10X files were loaded into R using the Read10X function of Seurat. A single Seurat object for all four patients was created using the CreateSeuratObject function of Seurat. At this step, genes not expressed in at least 3 cells at a threshold of a minimum of 1 UMI count were excluded from the analysis (min.cells = 3). The percentage of mitochondrial counts was then calculated for each cell using the PercentageFeatureSet function of Seurat. Cells were removed from the analysis if they had less than 400 detected UMI counts, expressed less than 100 or more than 6000 genes, and if their mitochondrial counts exceeded 10% of their total counts. This filtering resulted in retaining 28,726 cells and 26,439 genes.

#### Dataset integration

The standard Seurat version 3^64^ integration workflow was used to integrate the data from all patients, as described in Durante et al.^33^ First, the data from each patient was separated using the SplitObject function of Seurat. The raw UMI counts of each gene within each cell were normalized by the total number of UMI counts per cell, scaled to 10^4^ and natural-log-transformed after adding a pseudocount of 1 using the NormalizeData function of Seurat (normalization.method = “LogNormalize”; scale.factor = 10000). The top 5000 variable genes were then determined in each dataset using the variance-stabilizing transformation^64^ method of the FindVariableFeatures function of Seurat (selection.method = “vst”; nfeatures = 5000). To assemble all datasets, integration anchors (i.e. mutual nearest neighbors) across the four datasets were identified with the FindIntegrationAnchors function of Seurat after computing the first 30 dimensions of the canonical correlation analysis^65^ between each pair of datasets using the top 5000 variable genes shared between most datasets (reduction = “cca”; dims = 1:30; anchor.features = 5000; l2.norm = TRUE). The IntegrateData function of Seurat was then used to integrate all four datasets into a single Seurat object.

#### UMAP plot generation

To generate a UMAP^67,68^ plot, the integrated data was scaled and centered using the ScaleData function of Seurat and the first 30 principle components were computed using the RunPCA function of Seurat. These 30 principle components were then used as input to the RunUMAP function of Seurat (dims = 1:30). The UMAP plot was generated with the uwot R package version 0.1.8 implementation. The cosine metric was used to measure distance and find the nearest neighbors (metric = “cosine”), the number of neighbors used for the approximation of the manifold structure was set to 30 (n.neighbors = 30) and the effective minimum distance between embedded points was set to 0.3 (spread = 1; min.dist = 0.3). The clusters identified by Durante et al.^33^ were displayed on the UMAP plot (Figure 2B) and were used for subsequent analyses. The cluster corresponding to “Respiratory columnar cells” was renamed as “Respiratory early secretory cells” based on its expression of *SERPINB3* (see Supplementary Table 3 from Durante et al.^33^).

#### Gene expression analysis and plotting

All gene expression analyses were performed using the actual normalized UMI counts (not log-transformed) from the non-integrated assay of the Seurat object. However, the scales of the axes in the violin plots (Figure 2A, F and G) and the color gradient scales in the heatmap plots (Figure 2C) are in log 10 after adding a pseudocount of 1 to the normalized UMI counts. The axes in Figure 2E, H and I are not in log scale. *ACE2* and *TMPRSS2* coexpressing cells were identified as expressing at least 1 UMI count of both genes (Figure 2D). All violins in each violin plot have the same maximum width.

### Droplet-based single-nucleus RNA sequencing data analysis of human brain cells

#### Data download

The digital gene expression matrices of the single-nucleus Drop-seq dataset reported in Lake et al.^44^ were downloaded from the NCBI GEO database with the accession number GSE97930. This dataset consists in human brain nuclei collected from the frontal cortex, visual cortex and lateral cerebellar hemisphere from six individuals aged between 20 and 49 years (for more information, see Supplementary Table 1 from Lake et al.^44^). The data was analyzed in R version 3.5.0 using the pagoda2 R package (https://github.com/hms-dbmi/pagoda2) version 0.1.1 following the indications described in Lake et al.^44^ and using custom scripts. For simplification, the word “cells” will be used instead of “nuclei” in the following paragraphs.

#### Cell and gene filtering

All cells from the downloaded matrices had a cell type annotation and were therefore retained for further analysis. Genes shared between all 3 datasets (i.e. frontal cortex, visual cortex and lateral cerebellar hemisphere) were selected for downstream analysis. The final UMI count matrix, after combining all datasets, contained 35,289 cells and 31,302 genes.

#### Batch correction and data preparation

A new Pagoda2 object was created and the library identification of each cell was defined for batch correction using the plain model (see Supplementary Table 2 from Lake et al.^44^). Winsorization was used to clip the expression levels of each gene at the eleventh most extreme value (i.e. winsorizing the ten most extreme values; trim = 10). Batch-corrected gene expression levels were then normalized by the total number of counts per cell, scaled to 10^3^ and natural-log-transformed after adding a pseudocount of 1.

#### UMAP plot generation

The variance of genes were calculated and adjusted using the adjustVariance function of pagoda2, as described in Lake et al.^44^ (the smoothing term of the generalized additive model was set to gam.k = 10). The 2,000 overdispersed genes were then selected from the variance adjusted matrix and the 150 principle components were computed using the calculatePcaReduction function of pagoda2 (n.odgenes = 2000; nPcs = 150). These 150 principle components were then used to generate a UMAP plot with the getEmbedding function of pagoda2 (type = “PCA”; embeddingType = “UMAP”). As for the analysis of human olfactory epithelial cells, the uwot R package version 0.1.8 implementation was used to generate the UMAP plot. Also, the cosine metric was used to measure distance and find the nearest neighbors (distance = “cosine”), the number of neighbors used for the approximation of the manifold structure was set to 15 (n_neighbors = 15) and the effective minimum distance between embedded points was set to 0.1 (spread = 1; min_dist = 0.1).

#### Gene expression analysis and plotting

To plot the expression levels of *ACE2* and *TMPRSS2* on the UMAP projection in the form of a heatmap, the raw UMI counts of each gene within each cell were normalized by the total number of UMI counts per cell and scaled to 10^4^. Similar to Figure 2C, the color gradient scales in the heatmap plots (Supplementary Figure 4B) are in log_10_ after adding a pseudocount of 1 to the normalized UMI counts. In Supplementary Figure 4D and E, broader clusters were defined by separately grouping all excitatory neuron clusters, all inhibitory neuron clusters and all Purkinje neuron clusters.

### SMART-Seq v4 single-nucleus RNA sequencing data analysis of human brain cells

#### Data download

The digital gene expression matrix (matrix.csv), the *t*-SNE coordinates for each cell (tsne.csv) and the metadata table corresponding to the cells (metadata.csv) were downloaded from the Allen Brain Map cell types database, available online at https://portal.brain-map.org/atlases-and-data/rnaseq/human-multiple-cortical-areas-smart-seq. This dataset consists in human brain nuclei collected from three individuals and from eight brain regions: middle temporal gyrus (MTG), cingulate gyrus (CgG), primary visual cortex (V1C), primary auditory cortex (A1C), upper limb (ul) and lower limb (lm) regions of both primary motor cortex (M1) and primary somatosensory cortex (S1) (for more information, see metadata.csv from the website above). The data was analyzed in R version 3.5.0 using custom scripts. Like in the above paragraphs, the word “cells” will be used instead of “nuclei” in the following paragraphs.

#### Cell and gene filtering

Cells lacking a brain cell-type subclass annotation (subclass_label column from metadata.csv) and a *t*-SNE embedding were removed from the analysis, and all genes were retained for data normalization. The final expression matrix contained 47,432 cells and 50,281 genes.

#### Gene expression analysis and plotting

Prior to any gene expression plotting or analysis, the raw counts of each gene within each cell were normalized by the total number of counts per cell and scaled to 10^4^. Similar to Figure 2C, the color gradient scales in the heatmap plots (Figure 6B and C) are in log_10_ after adding a pseudocount of 1 to the normalized counts. However, the axes in Figure 6E, F and G are not in log scale. Also, as described above, *ACE2* and *TMPRSS2* coexpressing cells were identified as expressing at least 1 UMI count of both genes (Figure 6D).

### Data visualization and programming tools

All plots in Figures 1, 2 and 6, as well as in Supplementary Figure 4, were generated using ggplot2 version 3.3.0. Other R and Bioconductor packages used for data analysis and plotting include rtracklayer^69^ version 1.46.0, Matrix version 1.2.18, reshape2 version 1.4.3, data.table version 1.12.8, dplyr version 0.8.5, stringr version 1.4.0, ggrepel version 0.9.0, lemon version 0.4.4, extrafont version 0.17, viridis version 0.5.1, RColorBrewer version 1.1.2, patchwork version 1.0.0 and grid version 3.5.0.

### Mouse brain cell database mining

For mouse brain single-cell transcriptomes, the datasets of Zeisel et al.^41^ (Figure 5A and B) and Sauders et al.^42^ (Figure 5C) were analyzed from their respective websites: http://mousebrain.org/ and http://dropviz.org/.

### Immunohistochemistry and histology

#### Tissue fixation and preparation

All animal procedures were performed in accordance with the guidelines and regulations of the institution and of the state of Geneva. Mice (C57BL6, Charles River Laboratories, France) were deeply anesthetized with Pentobarbitalum natricum at 150mg/kg (Streuli Pharma) and perfused transcardially with 1x phosphate-buffered saline (PBS)-heparine 20’000 UI/l (Bichsel), pH 7.4. After flushing, mice were perfused with 10% neutral buffered Formalin. Tissues were extracted and fixed over night at 4°C and cryoprotected in 15% sucrose, 1x PBS over night at 4 °C followed by 30 % sucrose, 1x PBS over night at 4 °C. Tissues were embedded in optimal cutting temperature compound (OCT)(Carl Roth, ref. 6478.1) and cut on a cryostat in 16 μm sections that were mounted on Superfrost+ slides. Sections were conserved at −80°C until use. Human olfactory and respiratory epithelium was sampled from autopsy of a 95-year-old woman. By transcranial access, the cribriform plate with the superior part of the nasal cavities and the septum nasi were removed. The postero-lateral and postero-medial parts of the nasal mucosa were carefully dissected and fixed in 10% neutral buffered Formalin for 48 hours and bony particles were decalcified in EDTA for 24 hours. Paraffin embedding was done using a standard protocol using a LOGOS Microwave Hybrid Tissue Processor (Milestone). Paraffin blocks were cut at 3 μm thickness with a microtome and mounted on Superfrost slides. Before staining, cryosections were thawed and dried for 30 min under a ventilated hood. Sections were re-hydrated with 1x PBS for 5 min. Paraffin sections were deparaffinized by placing them for 10 min in xylene (Fluka), and then successively for 1 min each in ethanol 100%, 90%, 80%, 70% and H_2_O.

#### Alcian blue staining

For alcian blue staining, hydrated sections were stained for 30 min in alcian blue (Sigma, ref. 1.01647), acetic acid 3%, pH 2.5. Sections were washed for 2 min in running tap water and rinsed in distilled water. Sections were counterstained with 0.1% nuclear fast red, 5% aluminum sulfate solution (Merck Millipore, ref. 1.00121) for 5 min. Sections were washed for 1 min in running tap water and dehydrated. Sections were cleared in xylene and mounted with Eukitt (Sigma, ref. 03989) mounting medium.

#### Immunohistochemistry

For antigen retrieval, sections were immersed in 10mM Citric Acid, 0.05% Tween 20, pH 6.0 buffer previously heated to 95-100 °C, and incubated for 20 min. Sections were then cooled down to room temperature in 1x PBS. Section were pre-incubated for 30 min with 1x PBS containing 0.5% Triton X-100 and 5% FCS. Sections were incubated with primary antibodies (see details in Table 1) diluted in 1x PBS containing 0.5% Triton X-100 and 5% FCS over night at 4°C. Sections were washed 3×15 min with 1x PBS containing 0.5% Triton X-100. Sections were incubated with secondary antibodies diluted 1:800 in 1x PBS with 0.5% Triton X-100 and 5% FCS for 90 min at room temperature (see details in Table 2). Sections were again washed 3×15 min with 1x PBS containing 0.5% Triton X-100. Sections were counterstained with DAPI (1:5000) for 5 min, rinsed with 1x PBS and mounted with DABCO (Sigma) in glycerol. For all tissue samples and all conditions, control samples were processed simultaneously without applying primary antibodies.

**Table 1.**
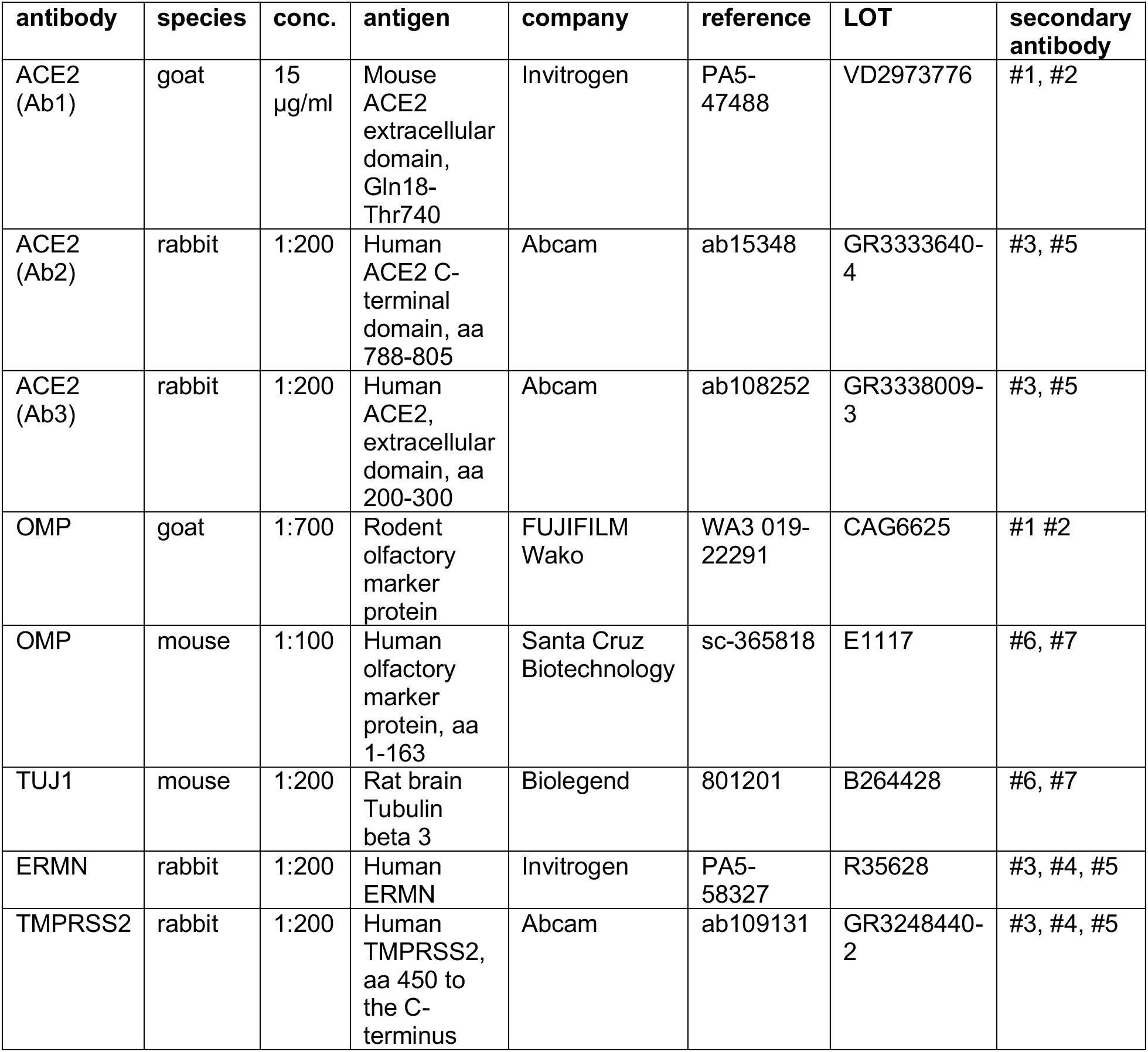
Primary antibodies

| antibody | species | conc. | antigen | company | reference | LOT | secondary antibody |
| --- | --- | --- | --- | --- | --- | --- | --- |
| ACE2 (Ab1) | goat | 15 µg/ml | Mouse ACE2 extracellular domain, Gln18-Thr740 | Invitrogen | PA5-47488 | VD2973776 | #1, #2 |
| ACE2 (Ab2) | rabbit | 1:200 | Human ACE2 C-terminal domain, aa 788-805 | Abcam | ab15348 | GR3333640-4 | #3, #5 |
| ACE2 (Ab3) | rabbit | 1:200 | Human ACE2, extracellular domain, aa 200-300 | Abcam | ab108252 | GR3338009-3 | #3, #5 |
| OMP | goat | 1:700 | Rodent olfactory marker protein | FUJIFILM Wako | WA3 019-22291 | CAG6625 | #1 #2 |
| OMP | mouse | 1:100 | Human olfactory marker protein, aa 1-163 | Santa Cruz Biotechnology | sc-365818 | E1117 | #6, #7 |
| TUJ1 | mouse | 1:200 | Rat brain<br>Tubulin<br>beta 3 | Biolegend | 801201 | B264428 | #6, #7 |
| ERMN | rabbit | 1:200 | Human<br>ERMN | Invitrogen | PA5-58327 | R35628 | #3, #4, #5 |
| TMPRSS2 | rabbit | 1:200 | Human<br>TMPRSS2,<br>aa 450 to<br>the C-terminus | Abcam | ab109131 | GR3248440-2 | #3, #4, #5 |

**Table 2.** Secondary antibodies

| # | antibody | species | company | reference |
| --- | --- | --- | --- | --- |
| 1 | anti-goat Alexa 555 | donkey | Invitrogen | A21432 |
| 2 | anti-goat Alexa 488 | donkey | Abcam | ab150133 |
| 3 | anti-rabbit Alexa 555 | donkey | Abcam | ab150074 |
| 4 | anti-rabbit Alexa 488 | donkey | Abcam | ab150073 |
| 5 | anti-rabbit Alexa 488 | goat | Life Technologies | A11034 |
| 6 | anti-mouse Alexa 488 | donkey | Abcam | ab150105 |
| 7 | anti-mouse Cy3 | goat | Life Technologies | A10521 |

#### Microscope imaging

Confocal imaging was performed with a Leica TCS SP8 using a PMT detector for DAPI fluorescence and HyD 5 detectors for Alexa488, Alexa555 and Cy3 fluorescence. Epifluorescence imaging was performed with a Leica DM5500 equipped with a DFC9000 GT monochrome camera. For both fluorescence microscopy techniques, illumination and detection was manually adjusted to optimize signal-to-noise ratio and minimize over-exposure, bleaching and light scattering. Brightfield imaging was performed with a Leica DM5500 equipped with a DMC2900 camera. Microscope automated control and image encoding was performed with the LAS X software version 3.4.2. Acquisition settings were matched in negative controls.

#### Microscope image editing

Maximum intensity projection were computed on the LAS X software for z-stack imaging. The following images are z-stack maximum intensity projections: Figure 1G, K, M, O, Q, Supplementary Figure 1A, B, Figure 4E-G. All images were exported as TIF files. For clarity, cropping and linear exposure adjustments were performed on Adobe Photoshop. Again, settings were matched in negative controls.

## Acknowledgements

We thank the iGE3 Genomics Platform at the University of Geneva for expert technical assistance during RNA-seq experiments. This research was supported by the University of Geneva and the Swiss National Science Foundation (grant numbers: 31003A_172878 to A.C. and 310030_189153 to I.R.).

## Author contributions

LF, JT, MB, DR, CK, and VP acquired and analyzed data. LF performed bulk, singlecell and single-nucleus RNA-seq data analysis. JT performed bulk RNA-seq data analysis, mouse brain cell database mining, and confocal microscopy. DR generated the human bulk RNA-seq dataset, MB performed immunohistochemical stainings and epifluorescence microscopy with help from CK and VP. KE, JL and BL collected human biopsies. LF, JT, MB, DR, AC and IR carried out the conceptualization and experimental design, AC and IR wrote the manuscript with comments from all the other authors.

## Supplementary Figures

**Supplementary Figure 1.**
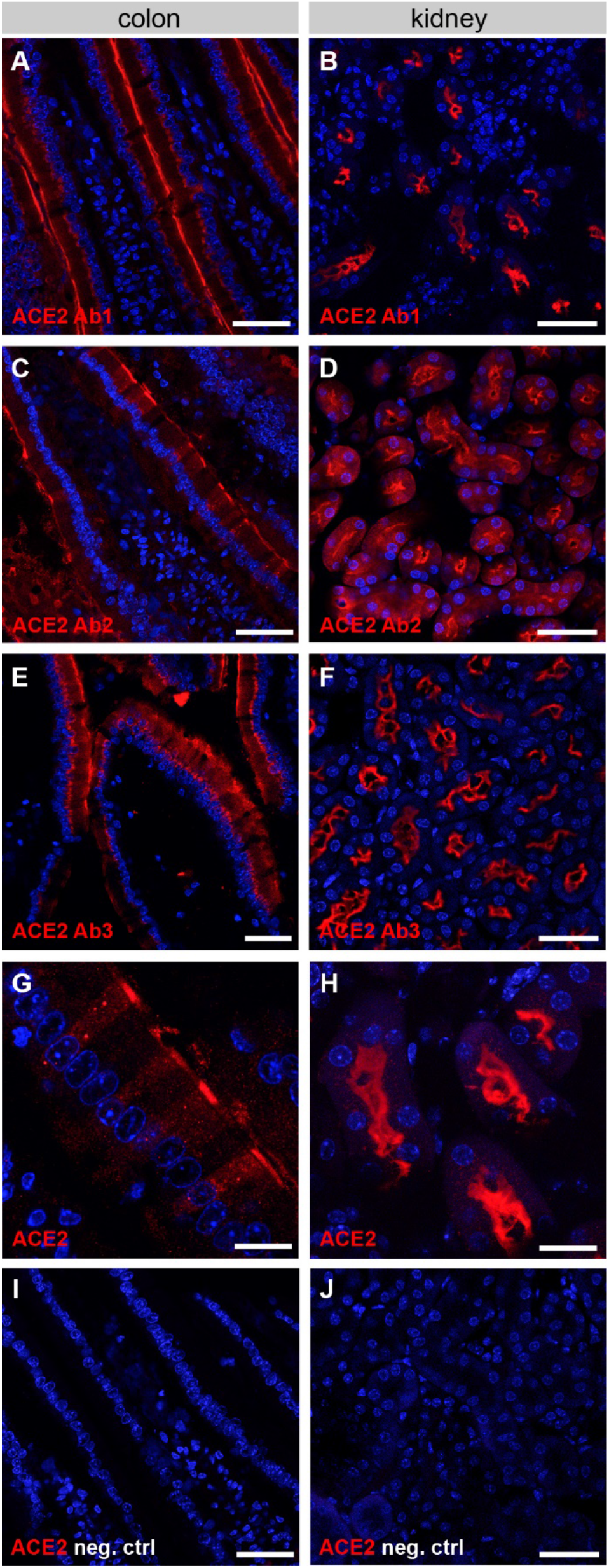
ACE2 expression in the mouse kidney and colon. (**A-H**) Evaluation of the three different anti-ACE2 antibodies. Immunostaining for ACE2 (red) on sections of mouse colon (left panels) and kidney (right panels). (**A,B**) ACE2 Ab1, polyclonal antibody against the extracellular domain of mouse ACE2. (**C,D**) ACE2 Ab2, polyclonal antibody against the intracellular domain of human ACE2. (**E**,**F**) ACE2 Ab3, monoclonal antibody against the extracellular domain of human ACE2. Counterstaining with DAPI (blue). Scale bar: 40 μm. (**G,H**) Higher magnification of mouse colon (**G**) and kidney (**H**) sections immunostained for ACE2 (red). Scale bar: 20 μm. (**I,J**) Negative control immunostainings on colon and kidney without primary antibody for ACE2. Scale bar: 40 μm.

**Supplementary Figure 2.**
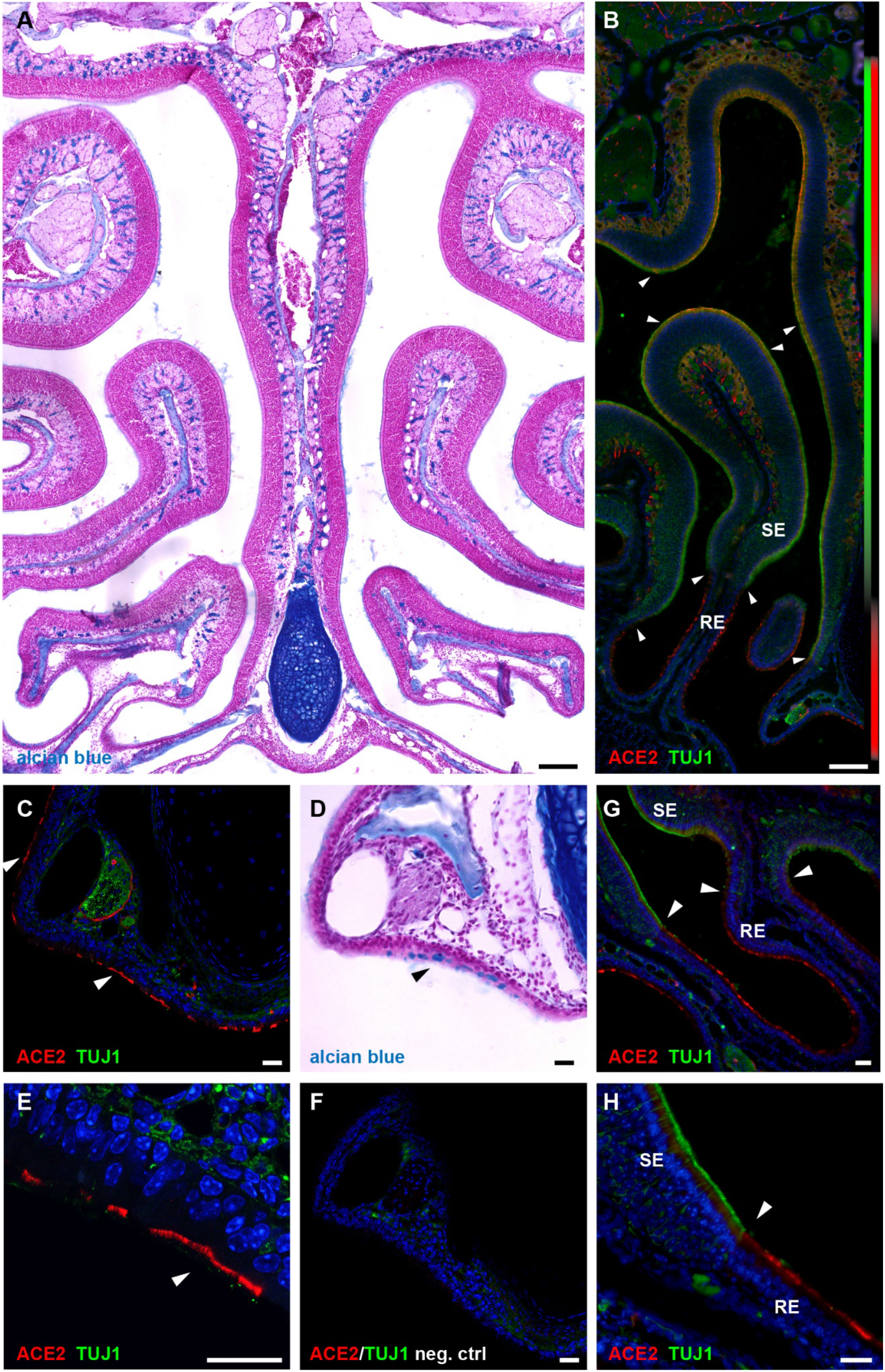
ACE2 expression in the mouse olfactory and respiratory epithelium. (**A**) Alcian blue staining on a coronal section of the mouse olfactory neuroepithelium. The entire sensory epithelium is devoid of goblet cells. Scale bar: 0.2 mm. (**B**) Immunostaining for ACE2 (red) and the neuronal marker TUJ1 (green) on a section of mouse nasal cavity and nasopharyngeal duct. White arrowheads indicate transitions from ACE2-positive to ACE2-negative zones. Colored bars on the right show expression zones of ACE2 (red) and TUJ1 (green). ACE2 is expressed in the dorsal olfactory epithelium and in the ventral respiratory epithelium. SE: sensory epithelium, RE: respiratory epithelium. Scale bar: 0.2 mm. (**C-H**) Evaluation of ACE2 expression in the mouse respiratory epithelium. (**C,E**) Immunostaining for ACE2 (red) and the neuronal marker TUJ1 (green) on sections of respiratory epithelium in the nasopharyngeal duct. (**D**) The same section with alcian blue staining, showing the presence of goblet cells. (**F**) Negative control immunostaining without primary antibodies. (**G,H**) Immunostaining for ACE2 (red) and the neuronal marker TUJ1 (green) on the ventral part of the nasal cavity, showing the transition from olfactory to respiratory epithelium (white arrowhead). Scale bar: 40 μm. (**H**) Higher magnification of the transition zone shown in (**G**). When not indicated, scale bars are 20 μm. All sections with immunostaining were counterstained with DAPI (blue).

**Supplementary Figure 3.**
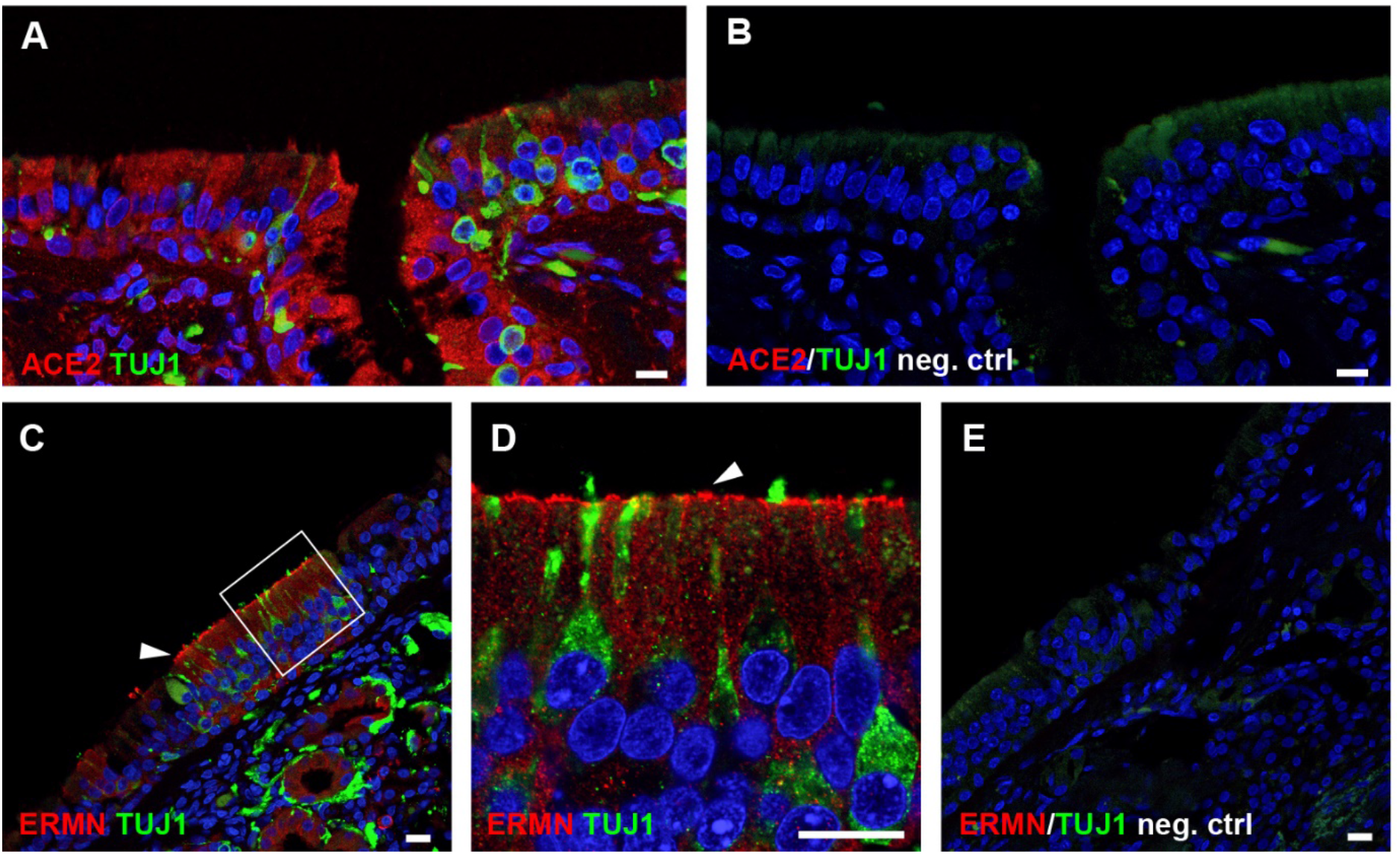
*ACE2* expression in the human olfactory neuroepithelium. (**A**) Section of human olfactory neuroepithelium immunostained for ACE2 (red) and for the neuronal marker TUJ1 (green). (**B**) Same region as in (**A**), immunostained without primary antibodies. (**C, D, E**) Section of human olfactory neuroepithelium immunostained for the sustentacular cell marker ERMN (red) and the neuronal marker TUJ1 (green). The white square in (**C**) highlights the region magnified in (**D**). ERMN is expressed in the apical portion of sustentacular cells (white arrowheads). (**E**) Same region as in (**D**), but without primary antibodies. All sections with immunostaining were counterstained with DAPI (blue). Scale bars: 10 μm.

**Supplementary Figure 4.**
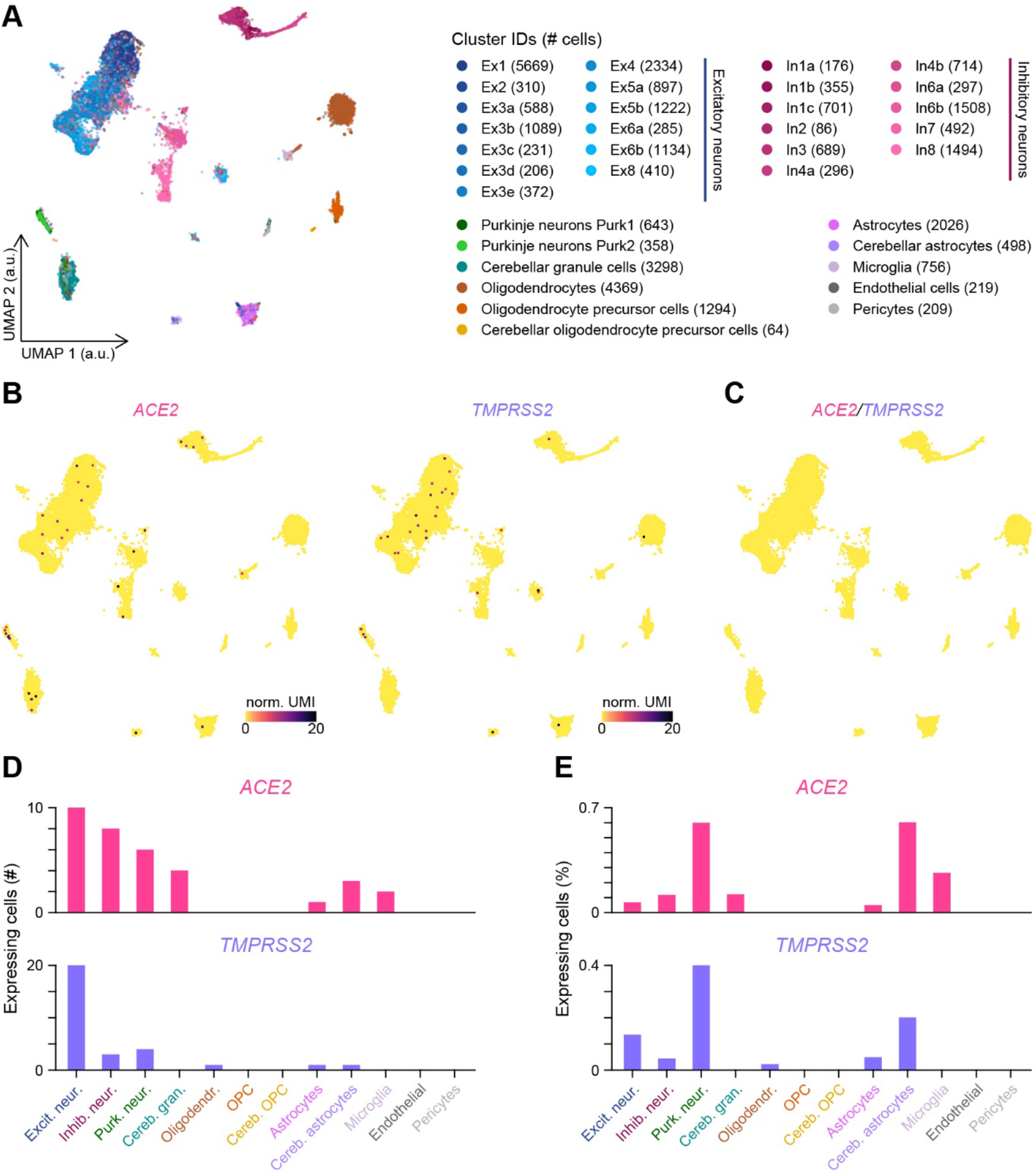
*ACE2* and *TMPRSS2* expression in the human brain. (**A**) Visualization of the clustering results reported in Lake et al.^44^ on a UMAP plot. a.u.: arbitrary units. (**B**) Normalized expression levels of *ACE2* and *TMPRSS2* shown on the UMAP plot. Color gradient scales are in log_10_. norm. UMI: normalized unique molecular identifier counts. (**C**) Absence of *ACE2* and *TMPRSS2* coexpressing cells highlighted on the UMAP plot. (**D,E**) Number (**D**) and percentage (**E**) of cells per cluster expressing *ACE2* or *TMPRSS2*. Combined results are shown for clusters with subdivisions (excitatory, inhibitory or Purkinje neuron clusters).

